# Inpainting as a Technique for Estimation of Missing Voxels in Chemical Shift Imaging

**DOI:** 10.1101/2020.02.17.952325

**Authors:** Angel Torrado-Carvajal, Daniel S. Albrecht, Jeungchan Lee, Ovidiu C. Andronesi, Eva-Maria Ratai, Vitaly Napadow, Marco L. Loggia

## Abstract

Issues with model fitting (i.e. suboptimal standard deviation, linewidth/full-width-at-half-maximum, and/or signal-to-noise ratio) in multi-voxel MRI spectroscopy, or chemical shift imaging (CSI), can result in the significant loss of usable voxels. A potential solution to minimize this problem is to estimate the value of unusable voxels by utilizing information from reliable voxels in the same image. We assessed an image restoration method called inpainting as a tool to restore unusable voxels and compared it with traditional interpolation methods (nearest neighbor, trilinear interpolation and tricubic interpolation). We applied these techniques to N-acetylaspartate (NAA) spectroscopy maps from a CSI dataset. Inpainting exhibited superior performance (lower normalized root-mean-square errors, NRMSE) compared to all other methods considered (p’s<0.001). Inpainting maintained its superiority whether the previously unusable voxels were randomly distributed or located in regions most commonly affected by voxel loss in real-world data.

**Clinical Relevance:** The presence of missing voxels can be problematic, particularly when data are analyzed in standard space, given that only voxels that are contributed to by all participants can be interrogated in these analyses. Inpainting is a promising approach for recovering unusable or missing voxels in voxelwise analyses, particularly in imaging modalities characterized by low SNR such as CSI.

## I. Introduction

The use of voxelwise analyses (e.g., to evaluate pathological alterations in a given patient group, or associations between imaging and clinical variables) is very widespread in brain imaging. Compared to region-of-interest (ROI) analyses, which require averaging the signal over multiple voxels across a predefined region, voxelwise analyses allow localizing effects of interest with greater spatial accuracy, often improving sensitivity to detect spatially restricted effects that may be diluted in ROI analyses [1-4].

On the other hand, the use of voxelwise analyses can come with a cost, in terms of signal-to-noise ratio (SNR). Chemical shift imaging (CSI), for instance, enables the performance of multi-voxel MRI spectroscopy, therefore conferring imagers the ability to interrogate metabolite levels across a larger portion of the brain, and with higher spatial resolution, compared to single voxel spectroscopy. However, the smaller size of the CSI voxels can lead to lower SNR and occasional poor spectral fits, ultimately leading to some percentage of voxels becoming unusable. While these unusable voxels can be more commonly expected in some parts of the brain (e.g., regions more likely affected by magnetic inhomogeneities, ventricles), in other parts their spatial distribution can be somewhat random and idiosyncratic across subjects. This becomes particularly problematic for analyses in standard space: once multiple CSI maps, each with a different pattern of unusable voxels, are brought to a standard space for higher level analyses, the resulting common standard space can be significantly reduced, and sometimes rendered overall unusable, given that only voxels that are contributed to by all participants can be interrogated in these analyses.

A potential solution to minimize this problem is to recover the missing data using interpolation methods, which allow the estimation of missing data points within the range of a discrete set of known data points [5]. While these methods are commonly used in medical imaging (e.g., when resampling volumes), their performance is often affected by errors in the fitting, resulting in discontinuities, granularity and unrealistic values [6, 7]. An alternative to traditional interpolation methods is provided by image inpainting [8, 9]. Inpainting, sometimes also called “fill in”, is a method that fills in missing or corrupted parts of an image by taking into account both its global and local features. To date, inpainting has not been broadly applied to brain imaging. The few applications published so far have assessed the ability of inpainting methods to alleviate the bias induced by multiple sclerosis lesions in morphometric estimates [10-12], or to correct artifacts created by metallic implants [13].

In this work, we explore the use of the inpainting approach as a way to restore poor CSI data. To do this, we compare the performance of inpainting against other, more commonly used interpolation methods. These results would serve as a proof of concept to set up the basis for future research on the use of these methods in brain imaging.

## II. Materials and Methods

### A. General Strategy

In this study, we evaluated the performance of inpainting and several common image interpolation methods on N-Acetyl aspartate (NAA) CSI maps. Two image degradation strategies were implemented. First, all images were corrupted in such a way as to cause the images to randomly lose 5% to 95% of voxels (“%loss”; in steps of 5%). The NAA maps used in these tests were raw, and not thresholded to meet typical quality control (QC) standards (see subsection II.C). These tests were performed once per %loss level for each of the available T1 or CSI images, each time with a different random distribution of missing voxels. Second, as low quality CSI voxels tend to be more expected in some areas of the brain (e.g., regions more likely affected by inhomogeneities or low SNR) rather than being randomly distributed, the same NAA maps were corrupted in the spatial distribution encountered in real-life scans, in order to validate the methods in a more realistic scenario. To this end, we identified NAA images from 5 subjects demonstrating a significant proportion (mean±SD: 28.3±8.8%; range 18.0-37.5%) of voxels not meeting a standard QC criterion (voxelwise SD<20%). From these subjects, we extracted masks representing “bad NAA voxels”, and affine registered each of them to the NAA maps from 10 subject images presenting less than 10% of missing voxels in the field of view (FOV) (7.4±1.9%, range 4.6-9.7%). The ability of the various methods to rescue the ground truth voxels (i.e., the voxels meeting QC standards) from the “good NAA maps”, after imposing onto them the missing voxels derived from the “bad NAA maps”, was assessed by computing the error of the estimates (i.e., the normalized root mean square error).

### B. Datasets

Eighteen healthy controls (HCs; mean age, 46.4±14.4 y; range, 23-66 y; 8 males / 10 females) were screened and enrolled in this study. Participants were excluded if they had any MR contraindications such as metallic implants, history of head trauma and/or claustrophobia, had a history of major medical or psychological disorders. A more detailed description of the studies and their criteria can be found in [14-16]. The research protocol was approved by the local Institutional Review Board and written informed consent was acquired from all participants.

MR imaging was performed in a 3T Siemens TIM Trio scanner (Siemens Healthineers) using an 8-channel head and neck coil and included an anatomical T1-weighted volume (MEMPRAGE; TR/TE1/TE2/TE3/TE4=2530/1.64/ 3.5/5.36/7.22ms, flip angle=7°, voxel size=1×1×1mm, acquisition matrix=280×280×208), and CSI (LASER excitation and stack-of-spiral 3D k-space encoding; TR/TE=1500/30ms, volume of interest VOI=100×80×50mm, FOV=240×240× 100, matrix 24×24×10, voxel size=10×10×10mm) [17].

### C. Data Processing

CSI raw spectra data were transferred to a workstation where metabolites were fitted with LCModel software [19] using a bias set simulated with GAMMA environment [20] for LASER excitation [21] with the same pulse modulation as the acquisition sequence. QC for spectral fitting included line width (full-width at half maximum FWHM) < 0.1 ppm (12Hz at 3T), SNR > 3, and Cramer Rao lower bounds < 20% (relative to the fitted value) [22]. Metabolic maps were reconstructed and registered to the anatomical images using a combination of MINC/FSL/Matlab tools [23].

### D. Inpainting

The aim of inpainting is to provide an estimation of missing/corrupted parts of an image so that it looks natural to the human eye. As opposed to other methods, inpainting takes into account global, in addition to local, properties of the image. In general, the missing regions in an image might be composed of structures and textures, and separating these properties in two different steps is essential, starting by first recovering the structures and then recovering the texture [24].

Several inpainting approaches were developed in computer vision (variational image inpainting, texture synthesis, image completion, etc) [9]. For the purpose of its application to brain imaging, we have assessed a Matlab (Natick, MA) implementation based on a penalized least squares regression method that allows restoring missing data by means of the discrete cosine transform (DCT-PLS) [25, 26]. This method uses nearest neighbor interpolation to obtain a rough initial guess. Then, it introduces the texture information by deriving a statistical model that expresses the data in terms of a sum of cosine functions oscillating at different frequencies. Reference [25] contains detailed mathematical details of this method.

### E. Multivariate Interpolation Methods

As comparators, we have used three different multivariate interpolation methods commonly used in medical imaging: nearest neighbor, trilinear and tricubic interpolation [5]. The function to be interpolated is known in the surrounding voxels (*i*_*x*_, *i*_*y*_, *i*_*z*_) and the interpolation problem consists of approximating the value of an unknown voxel (*j*_*x*_, *j*_*y*_, *j*_*z*_). These methods were also implemented in Matlab.

### F. Analysis

Quantitative performance of the different methods was assessed by comparing the normalized root mean square error (NRMSE) computed between ground truth images and interpolated/inpainted images, using a repeated measures analysis of the variance (ANOVA) to evaluate the effect of method (inpainting, nearest neighbor interpolation, trilinear interpolation, tricubic interpolation), %loss (from 5 to 90%), and their interaction. Bonferroni-corrected T-tests were computed as follow-up tests, comparing inpainting against the other methods for each %loss level. Bonferroni correction was achieved by multiplying the *p* value by the number of %loss levels (*N* = 19); e.g., *p*_*corr*_ = 0.05 corresponds to *p*_*uncorr*_ = 0.00263. In order to assess the different methods in the real-life loss distribution scenario, we used a one-way ANOVA on ranks (Kruskal-Wallis), followed by paired-samples Wilcoxon signed rank tests to assess if the performance of inpainting was significantly different from the rest of the methods.

## III. Results

Overall, different methods filled-in missing pixels/voxels with acceptable results. However, inpainting outperformed the other methods. **Figure 1** shows the assessment on NAA CSI images. Despite the inherent low resolution of these images, we can appreciate the noise introduced by the interpolation methods, with inpainting being the method that maintained a voxel distribution that was the most similar to that of the original images (**Figure 1B**).

**Fig. 1.**
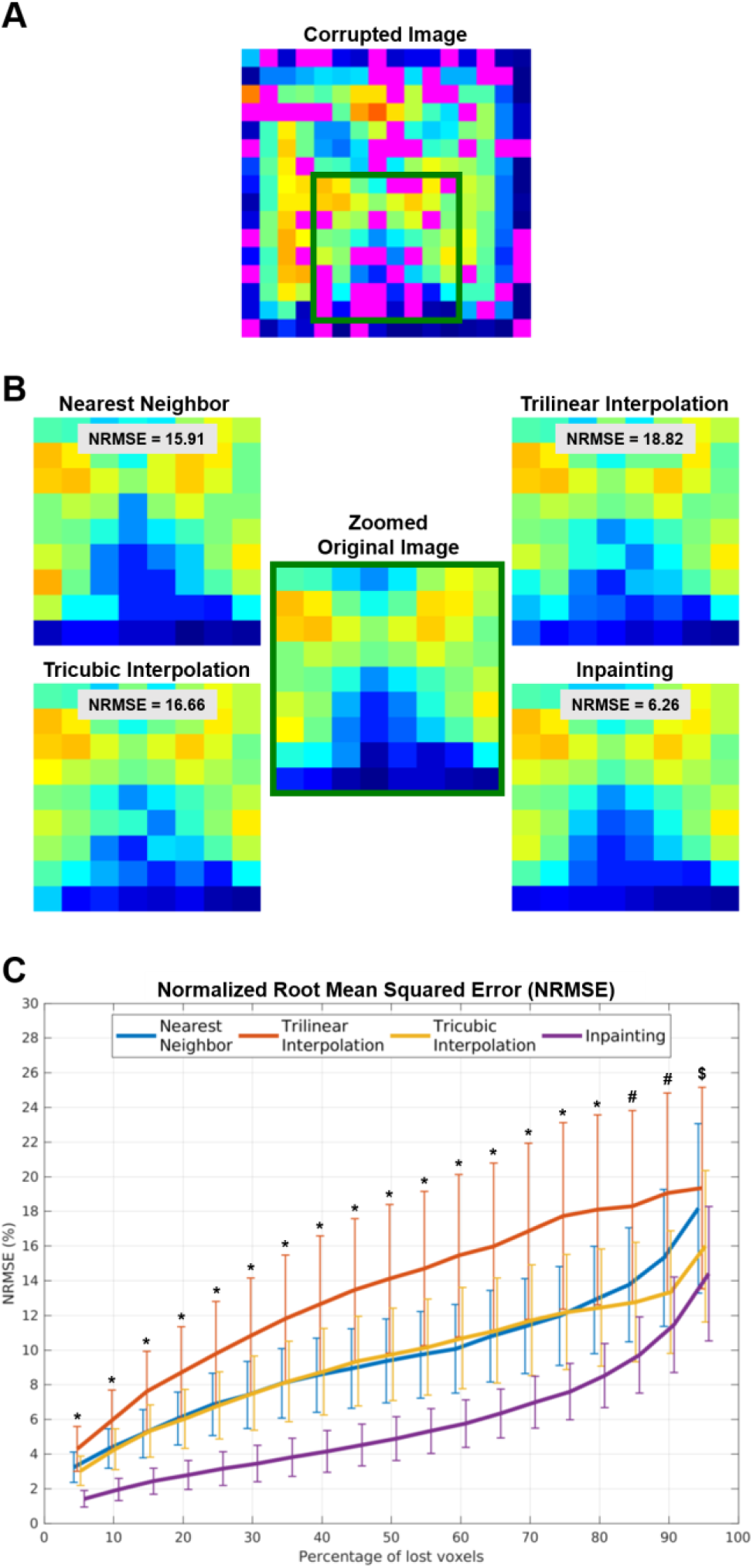
(A) NAA CSI image showing 50% of corrupted voxels (in magenta). (B) Zoomed view displaying details of the images restored using the various methods. (C) Normalized Root Mean Square Error (NRMSE) for the different methods, across different percentages of voxels loss. * *p*’*s* < 0.001 when directly comparing inpainting against all other methods in pairwise post-hoc analysis; # *p*′*s* < 0.05 only when directly comparing inpainting against nearest neighbor and trilinear interpolation. $ no statistical differences between inpainting and any of the other methods.

The quantitative assessment of the different methods shows that inpainting produced an average NRMSE of less than 5% even when with 50% missing voxels, whereas the other techniques demonstrated approximately twice (nearest neighbor, tricubic interpolation) or three times (trilinear interpolation) that value at that %loss level (**Figure 1C**). The ANOVA revealed a statistically significant effect of method (*F*_3,51_ = 99.9, *p* < 0.001) and percentage of lost voxels (%loss; CSI: *F*_18,306_ = 243.1, *p* < 0.001). We also observed a statistically significant method * %loss interaction (*F*_54,918_ = 76.4, *p*′*s* < 0.001), and the decomposition of the interaction using post-hoc pairwise comparisons revealed that inpainting outperformed all other methods at all %loss levels up to 80%loss (*p*’*s* < 0.001). Inpainting also outperformed nearest neighbor and trilinear interpolation for %loss levels between 85%loss and 90%loss (*p*’*s* < 0.05). No statistical differences were found for a 95%loss level.

In order to validate our methods in a more realistic scenario, the same NAA maps were corrupted in the same spatial distribution encountered in a dataset from actual CSI scans (**Figure 2**). We extracted masks representing “bad NAA voxels” (SD > 20%) for those subject images losing more than 15% of voxels in the FOV and registered those masks to images presenting an excellent NAA amplitude map. **Figure 2D** shows the histograms for the NRMSE values computed for the different methods. The Kruskall Wallis test revealed a statistically significant effect of method (*p* < 0.001). Follow-up post-hoc Wilcoxon signed rank tests showed significant differences in the inpainting NRMSE, as compared to the nearest neighbor, the trilinear and the tricubic interpolation methods (*ps* < 0.001). These results indicate that, once again, inpainting performed better than other methods, even when the voxels to be recovered had a spatial distribution consistent with that of actual CSI low-quality data.

**Fig. 2.**
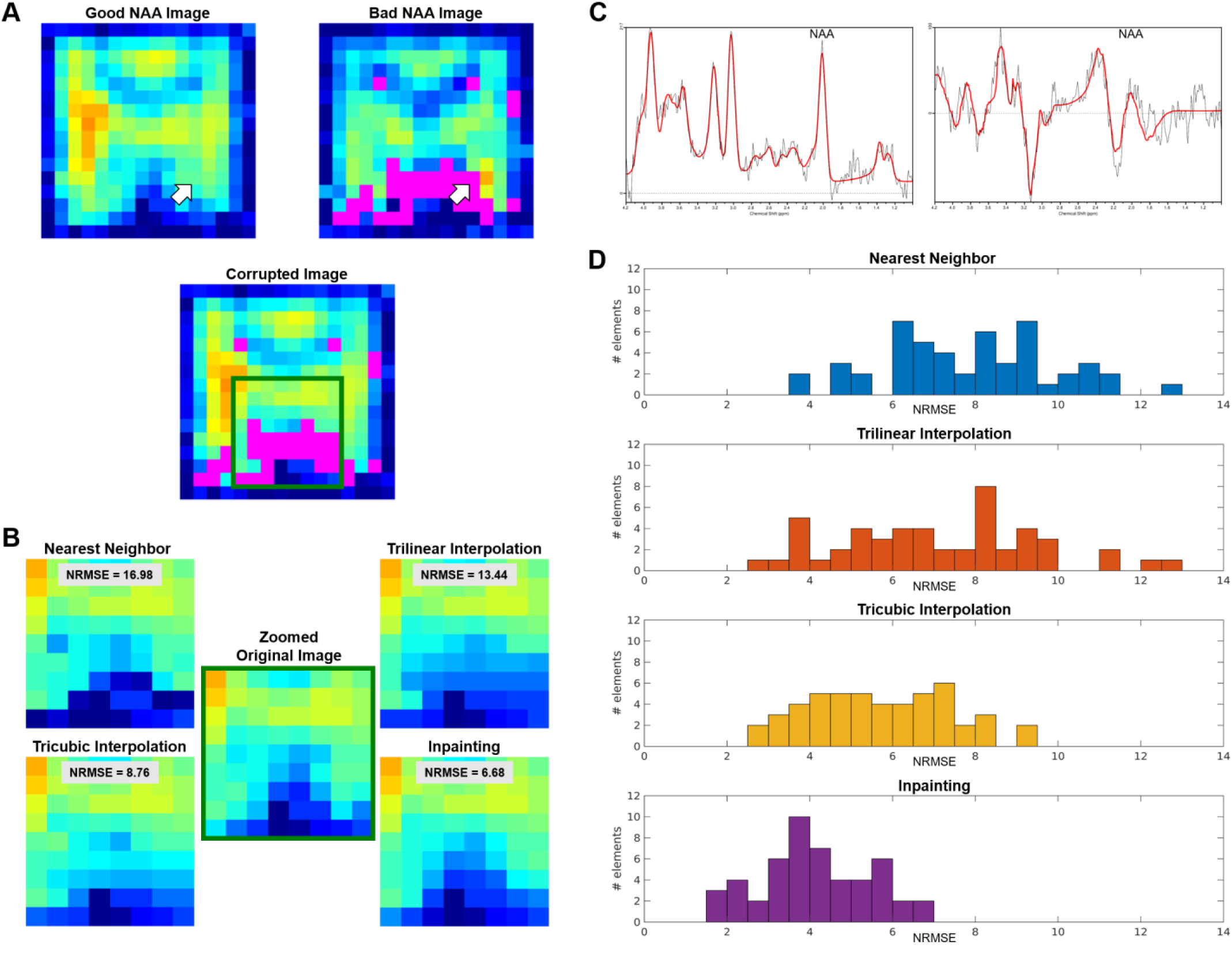
(A) Examples of a good NAA CSI image (left) and a bad NAA CSI image with a significant proportion of voxels (in magenta) not meeting QC threshold (right). Below is the result of imposing the mask from the bad image onto the good image. (B) Zoomed view displaying details of the images restored using the various methods and their performance to maintain the ground truth distribution. (C) Spectra and fittings corresponding to a good voxel (left) and a bad voxel (right) pointed by the white arrows in panel A. (D) Histogram of NRMSE distribution for the four different methods. NRMSE are binned in 0.5 wide bins.

## IV. Discussion

For a CSI voxel estimator to be considered viable, it needs to meet several minimum QC criteria including, but not limited to, the standard deviation of the estimators, the linewidth or FWHM of their spectrum, and/or the SNR in a given location. When a voxel does not meet these criteria, it is typically excluded. Our study evaluated the use of the inpainting approach as a tool to estimate values of previously unusable or missing voxels, such as in the case of CSI voxels not meeting minimum QC criteria.

Here, we directly compared the results of inpainting and other techniques, by corrupting data and evaluating how well each method recovered the original values. Overall, our results show that the application of image inpainting methods yields significantly better results compared to traditional interpolation methods. The performance of inpainting maintained its superiority compared to the other methods, whether the voxels to be estimated were randomly distributed, or located in regions genuinely prone to voxel loss in an actual CSI dataset (i.e., as estimated from real maps).

Of note, because brain NAA CSI datasets generally exhibit high SNR and low numbers of poorly fit voxels, the use of voxel-recovery methods might be even more relevant for other metabolites (e.g., glutamate/glutamine, myoinositol, etc). However, in this study we purposefully chose to apply our analyses on NAA maps because the relatively low percentage of poor quality voxels allowed us to compare the results of the various methods against a “ground truth”. Our study shows that inpainting could provide satisfactory results when applied to CSI data, despite their relatively small number of voxels, and inherent low spatial resolution.

Several limitations should be pointed out. First, our approach was applied on a relatively small number of datasets. Increasing the sample size would be desirable to report more accurate results. Second, the validation was performed using data from healthy controls subjects. Thus, in order to further evaluate the reliability of inpainting as a tool for data restoration, this method should be further validated on images with lesions, (i.e., tumors), and other abnormalities. Third, we applied these methods based on the premise that when a voxel does not meet certain QC criteria it is discarded (i.e. treated as having zero information). However, even low-SNR image regions do in fact contain information and we could initialize these methods based on the actual data instead of treating them as missing voxels.

In conclusion, our exploratory study demonstrated the use of an image restoration technique called inpainting to help recover poorly fitted CSI voxels with good accuracy. Future studies will need to evaluate to which extent the application of inpainting methods would be beneficial when applied to other imaging modalities as well.

